# Mass Spectrometry Imaging of Cocaine, Drug Removal and Methylphenidate Alter Phospholipids in Drosophila Brain

**DOI:** 10.1101/2020.01.08.899401

**Authors:** Mai H. Philipsen, Nhu T. N. Phan, John S. Fletcher, Andrew G. Ewing

## Abstract

Cocaine dependence displays a broad impairment in cognitive performance including attention, learning and memory. To obtain a better understanding of the action of cocaine in the nervous system, and the relation between phospholipids and memory, we have investigated whether phospholipids recover in the brain following cocaine removal using the fly model, *Drosophila*. In addition, the effects of methylphenidate, a substitute medication for cocaine dependence, on fly brain lipids after cocaine abuse are also determined to see if it can rescue the lipid changes caused by cocaine. Time of flight secondary ion mass spectrometry with a (CO_2_)_6000+_ gas cluster ion beam was used to detect intact phospholipids. We show that cocaine has persistent effects, both increasing and decreasing the levels of specific phosphatidylethanolamines and phosphatidylinositols. These changes remain after cocaine withdrawal and are not rescued by methylphenidate. Cocaine is again shown to generally increase the levels of phosphatidylcholines in the fly brain; however, after drug withdrawal the abundance of these lipids returns to the original level and methylphenidate treatment of the flies following cocaine exposure enhances the reversal of the lipid level reducing them below the original control. The study provides insight into the molecular effects of cocaine and methylphenidate on brain lipids. We suggest that phosphatidylcholine could be a potential target for the treatment of cocaine abuse, as well as be a significant hallmark of cognition and memory loss with cocaine.

## Introduction

Cocaine addiction is a worldwide serious health problem. One of the primary mechanisms of cocaine action is its ability to block presynaptic dopamine transporters which results in an increase in dopamine levels in the synaptic cleft resulting in euphoria and reinforcement of the administration of cocaine. Once the dopamine transporters are blocked by cocaine, most of the dopamine in the synaptic cleft cannot be recycled via re-uptake (1). Consequently, long-term repeated cocaine exposure inhibits dopamine transporters and consequently causes depletion of dopamine concentration and dysregulation of dopamine signaling in the brain (2). Neurochemical actions of cocaine also lead to an increase or a decrease in intracellular activities, especially the synthesis and degradation of phospholipids, in the reward-related brain regions which are associated with learning and memory (3). These changes in the brain might contribute to the deficit of brain functions observed including memory and cognition loss. Intriguingly, there is increasing literature showing that the symptoms of cognitive impairment are persistent long after cocaine withdrawal (4). However, there is still not effective medication available for cocaine abuse to date. One of the pharmacotherapeutic approaches for cocaine treatment is the agonist replacement therapy in which methylphenidate (MPH), for instance, can be used as a potential substitution for cocaine to reduce the cycle of compulsive use of cocaine. MPH also inhibits dopamine re-uptake which induces drug addiction; however, its clearance in the brain is slower and therefore the abuse potential is lower compared to cocaine. MPH has been showed to exert opposite effects to cocaine on cognitive function, noticeably it helps improve focus and memory for attention deficit hyper-activity disorder (ADHD) patients. Therefore, MPH might be a good candidate for treatment of cognition deficit caused by cocaine abuse.

Phospholipids are highly abundant in the brain and play important roles in brain function, particularly phospholipids regulate membrane trafficking, membrane fusion during exocytosis and endocytosis, regulate neurotransmitter receptors for neuronal signaling and transmission, and supply energy for brain activity (5). Phospholipids are composed of different groups with specific types of headgroups and tails, which determines their physical and chemical properties as well as their shapes and morphologies. These properties in turn result in different effects of the lipids on the activities of other neuronal molecules, for example, alteration of presynaptic protein interactions, activity of ion channels, and membrane curvature and fluidity (6). Phosphatidylcholines (PCs) and cholesterol have been shown to diminish in the cellular membrane of PC12 cells following exposure to cisplatin, and this is suggested as a molecular mechanism underlying the change of neurotransmitter release by this drug (7). In addition, various neurological disorders and brain diseases such as Alzheimer’s, ischemic stroke, ADHD, and schizophrenia (8–12) have been shown to result in alteration of brain lipids regarding their localization, abundance, and metabolism. MPH, as a common drug for ADHD patients, has been shown to change the lipid structure of the *Drosophila* melanogaster brain, especially by decreasing the abundance of cylindrical shaped phosphatidylcholines, and increasing the amount of conical shaped phosphatidylethanolamine (PEs) and phosphatidylinositols (PIs) (13). Furthermore, from our previous study comparing the effects of cocaine and MPH on the lipid structure of the fly brain (14), it is apparent that these two drugs lead to opposite changes in differing phospholipid levels. In that work, we hypothesized this as a possible mechanism for the opposite effects of cocaine and MPH on learning and memory.

Mass spectrometry imaging (MSI) is a powerful tool for spatial interrogation of altered lipid biochemistry (15). One of the MSI techniques, time-of-flight secondary ion mass spectrometry (ToF-SIMS), has been increasingly used in bioimaging to explore the lipid molecular architecture of cells and tissues (16). Given the high complexity and heterogeneity of the brain, it is essential to obtain detailed spatial organization in order to gain the insight of the relations between the molecular structure and function of the brain. ToF-SIMS offers the advantage that various molecules can be analyzed in parallel to obtain rich spatial molecular information inside the brain. The use of a high energy gas cluster ion beam (GCIB) with ToF-SIMS makes it particularly well suited for imaging intact phospholipids owing to the reduced fragmentation that occurs during sputtering by large primary ion clusters which therefore preserves the molecular ions (17, 18).

In this paper, we use ToF-SIMS imaging to test the hypothesis that the withdrawal of cocaine or addition of MPH after drug in *Drosophila* brain reverses the dramatic changes in the brain lipid composition caused by the exposure to cocaine. We have investigated if this strategy can be used to rescue the adverse effects of cocaine on lipid composition and brain functions. Interestingly, we find that cocaine removal as well as MPH treatment after cocaine only partially reverse the phospholipid changes in the brain caused by cocaine. This treatment seems to amplify the effects of cocaine removal on PCs and partially on TAGs, but neither cocaine removal nor MPH alter the levels of PE and PI species determined to change after cocaine. These seem to be more permanent changes that cannot be easily rescued.

## Materials and Methods

### Chemicals

All chemicals were purchased from Sigma Aldrich, Sweden.

### Drosophila preparation

Wide type *Drosophila* strain, Canton-S, was maintained in standard cornmeal food at room temperature (23- 26oC) with a 12 h light/dark cycle. Three to four-day-old male flies were selected and fed with yeast paste food supplemented with cocaine to get final concentration to 15 mM for 3 days. Cocaine-fed flies were divided into 3 groups: the first group had continued cocaine feeding for another 3 days; the second group, the so called cocaine removal group, was subsequently fed with normal yeast paste food without cocaine for 3 days; and the third group was orally administered with food containing 10mM MPH for 3 days after cocaine. After drug treatment, flies from different groups were placed sequentially on the same fly collars (4M Instrument & Tool LLC) and embedded in 10% gelatin (13). The gelatin molds were frozen at −20°C and then in liquid nitrogen where fly head-embedded gelatin was detached from the collar. The fly head-embedded gelatin was subsequently cut with 12 μm thickness using a cryo-microtome at −20°C (Leica CM1520). Sections were thaw-mounted on indium tin oxide coated glass and freeze-dried overnight in the ToF-SIMS instrument for further analysis.

### ToF-SIMS analysis - J105

ToF-SIMS analysis was performed using the J105-*3D Chemical Imager* (Ionoptika Ltd., UK). The principle operation of this instrument has been described in detail elsewhere (19). In our experiments, a GCIB of 40 keV (CO_2_)_6000+_ was used as a primary ion beam to sputter the sample surface in positive and negative ion modes. High energy clusters help enhance the primary and secondary ion yields for intact lipid signals in biological samples and allow detection of high mass species both on the sample surface and in the subsurface (18). The spectra were acquired over a mass range of *m/z* 100-1000. The image area was 1000 × 1000 μm2 with 128 × 128 pixels for statistical analysis, or with 256 × 256 pixels for imaging with higher spatial resolution, and the primary ion beam was 17 pA, which resulted in the total ion dose density of 6.1 × 10_12_ ions/cm_2_ and 1.7 × 10_13_ ions/cm_2_, respectively. The samples were analyzed in the freeze-dried condition at room temperature.

### Data analysis

SIMCA (Umetrix, Sweden) was used to perform principal components analysis (PCA) on ToF-SIMS spectra which were extracted by selecting the region of interest - the central area of the fly brain - from ToF-SIMS images. All spectra were normalized to total number of pixels and total ion intensity of the region of interest to compensate for the variations from ToF-SIMS measurements. PCA was then applied to this data set after Pareto scaling. Pareto scaling used square root of standard deviation as a scaling factor to reduce the dynamic range in the data and emphasize weak peaks that might be more relevant to biological information.

## Results

### ToF-SIMS for identifying phospholipid distributions on drug treated fly brain

ToF-SIMS is a powerful approach for studying lipidomic profiles and distributions in biological tissues. Due to the use of the relatively new GCIB technology, ion sources such as Ar_4000+_ and (CO_2_)_6000+_ provide greatly enhanced sputter yields of high mass molecular ions. Thus, a variety of intact phospholipids can be detected in fly brain sections. In order to evaluate the long-term effect of cocaine withdrawal or follow up MPH treatment on the phospholipid structure of the brain, we used ToF-SIMS with a (CO_2_)_6000+_ GCIB to image phospholipid localization in fly brain sections after the different drug treatments. One group of flies was treated with cocaine for 3 days and then fed with normal yeast-based food for 3 more days. Another set of cocaine-treated flies was subsequently treated by oral administration of MPH for 3 days. Comparison of the ion images of individual phospholipid species indicates a clear change in their distribution after cocaine removal or MPH treatment. In the positive ion mode, in the control flies, PCs, for instance PC (36:1), are mostly located in the central brain and some parts of optical lobe, especially the lobula complex (Fig. 1A). Interestingly, the PC species spread out to the medulla region of the optical lobe in the cocaine-treated brain, whereas after cocaine removal, their distributions return to those observed in the control group. In addition, the regional intensity of PC species increases in abundance mostly in the central brain after cocaine, then decreases to control levels after cocaine removal, but shows even lower abundance and is more focused in the central brain region of cocaine-fed flies when followed by MPH administration during cocaine removal. In the negative ion mode, the data show that PE and PI species are distributed more evenly across the whole control brain, while showing slightly higher intensity in the central brain and optical lobe regions. After cocaine administration, there are different trends in the distributions. For example, an enhancement of PE (34:1) intensity is observed in the cocaine-treated flies compared to the control flies. This is more or less uniform across the brain, but appears a little stronger in the optical lobes. This increase remains at the same level after the flies are removed from cocaine exposure for 3 days, and the abundance is even more pronounced after being treated by MPH following the cocaine (Fig. 1B). In contrast to PE (34:1), unsaturated PEs and PIs with 36 carbons in the fatty acid chains are decreased in the central brain and the optical lobe in the cocaine-fed fly brains compared to control flies. These also remain unchanged after cocaine removal, or after MPH treatment (Fig. 1C, D), although PI, such as PI (36:3), appears to recover slightly in the brains treated with MPH during cocaine recovery (semi quantitative data are given below). Overall, the data show that cocaine removal and MPH treatment after cocaine exposure reverse the effects of cocaine on the PC distributions in the fly brain, whereas there is no significant reversal in the changes in the perturbed PE and PI lipids caused by cocaine when it is then removed or the flies are exposed to MPH during the recovery time.

**Fig. 1.**
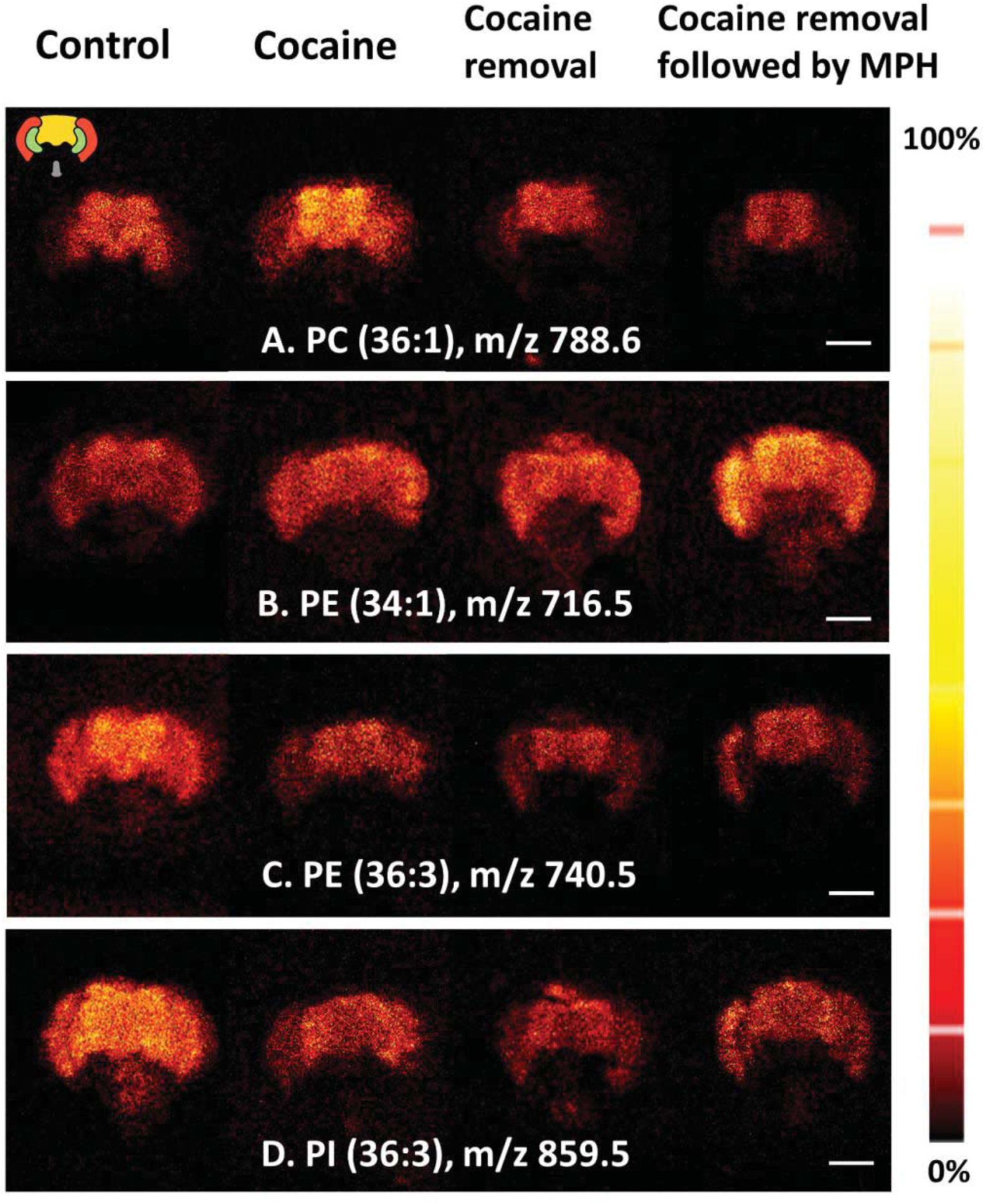
ToF-SIMS ion images show accumulation of phospholipids in fly head analyzed by ToF-SIMS with a 40 keV (CO_2_)_6000+_ GCIB. (A) PC(36:1), [M+H]+, in positive ion mode, (B, C) PE(34:1) and PE(36:3), [M-H]-, in negative ion mode, (D) PI(36:3), [M-H]-, in negative ion mode. Primary ion dose density 1.7 × 10_13_ ions/cm_2_. Image area is 1000 × 1000 μm2 and 256 × 256 pixels. Scale bar is 200 μm. Signal intensity is displayed on a color thermal scale on the right panel. Symbolic figure on the top left show the orientation of the fly head section including two red parts are medulla of optical lobes, two green parts are optical lobula, the central brain is the yellow part in the middle, and the grey part is proboscis.

### PC abundance in fly brains is rescued from cocaine effects by cocaine withdrawal and MPH treatment

Previous work showed that the effects of cocaine on the fly brain were most prevalent in the alteration of phospholipid compositions mainly in the central brain area, but no significant changes were observed in the optical regions and proboscis (14). In the current study, all the peaks in the spectra have been collected from the central regions of the fly brain and to elucidate the chemical changes, PCA was performed on ToF-SIMS spectra. These were normalized to the number of selected pixels and total ion counts. The scores plot of PCA from the extracted data clearly distinguishes three different groups along the second and third principal components.

Fig. 2A shows a clear separation in principal component 3 for the control and cocaine removal groups from the drug-treated groups. In Fig. 2B, the loadings plot of principal component 3 is included showing which peaks contribute most to the separation observed in the scores plots of principal component 3. The separation is mainly caused by peaks in the intact lipid region (e.g. triacylglycerol (TAG) and PC species). The assignment of peaks, which show significant alterations in principal component 3, can be found in Table S1. From Fig. 2B, PC species, for example PC (36:1) at *m/z* 788.6, PC (34:1)+K at *m/z* 798.5, and PC (36:3)+K at *m/z* 822.5, are more abundant in the purely cocaine-fed group and the group fed cocaine followed by MPH treatment. Likewise, other fragments of PC salt adducts such as PC (34:2)+Na-TMA at *m/z* 721.5, and PC (36:2)+Na-TMA *at m/z* 749.5, where TMA (59 Da) is the trimethylamine group at the end of the PC head group, display more intense intensity in cocaine-fed flies and cocaine-fed flies followed by MPH treatment. In contrast, elevations of the levels of TAG salt adducts, such as TAG (48:1)+Na at *m/z* 827.7, TAG (48:1)+K at *m/z* 843.7, and TAG (50:1)+K at *m/z* 871.7, are observed in control brains and those after cocaine removal. The diacylglycerol (DAG) peaks in the mass range m/z 400-650 are probably the fragments from TAGs and other lipids, hence we did not attempt to identify these species. In addition, use of principal component 2 separates the cocaine-fed group from the cocaine-fed followed by MPH group (Fig. S1). Specifically, the control, cocaine-fed, and cocaine removal groups have higher abundances of PCs and their salt adducts, such as PC (34:1) at *m/z* 760.6, PC (36:2)+K at *m/z* 824.6 and PC (36:2)+K-TMA at *m/z* 765.5, compared to the cocaine-fed group followed by MPH treatment. The signal level for the PC head group at *m/z* 184.1, which is correlated with PC species, is also decreased in cocaine-treated flies followed by MPH during the cocaine removal time. In contrast, the flies fed cocaine followed by MPH treatment during the cocaine removal time have spectra that are more dominant in the content of TAGs for example TAG (48:1)+K at *m/z* 843.7, TAG (50:2)+Na at *m/z* 853.7 and TAG (52:2)+Na at *m/z* 881.8.

**Fig. 2.**
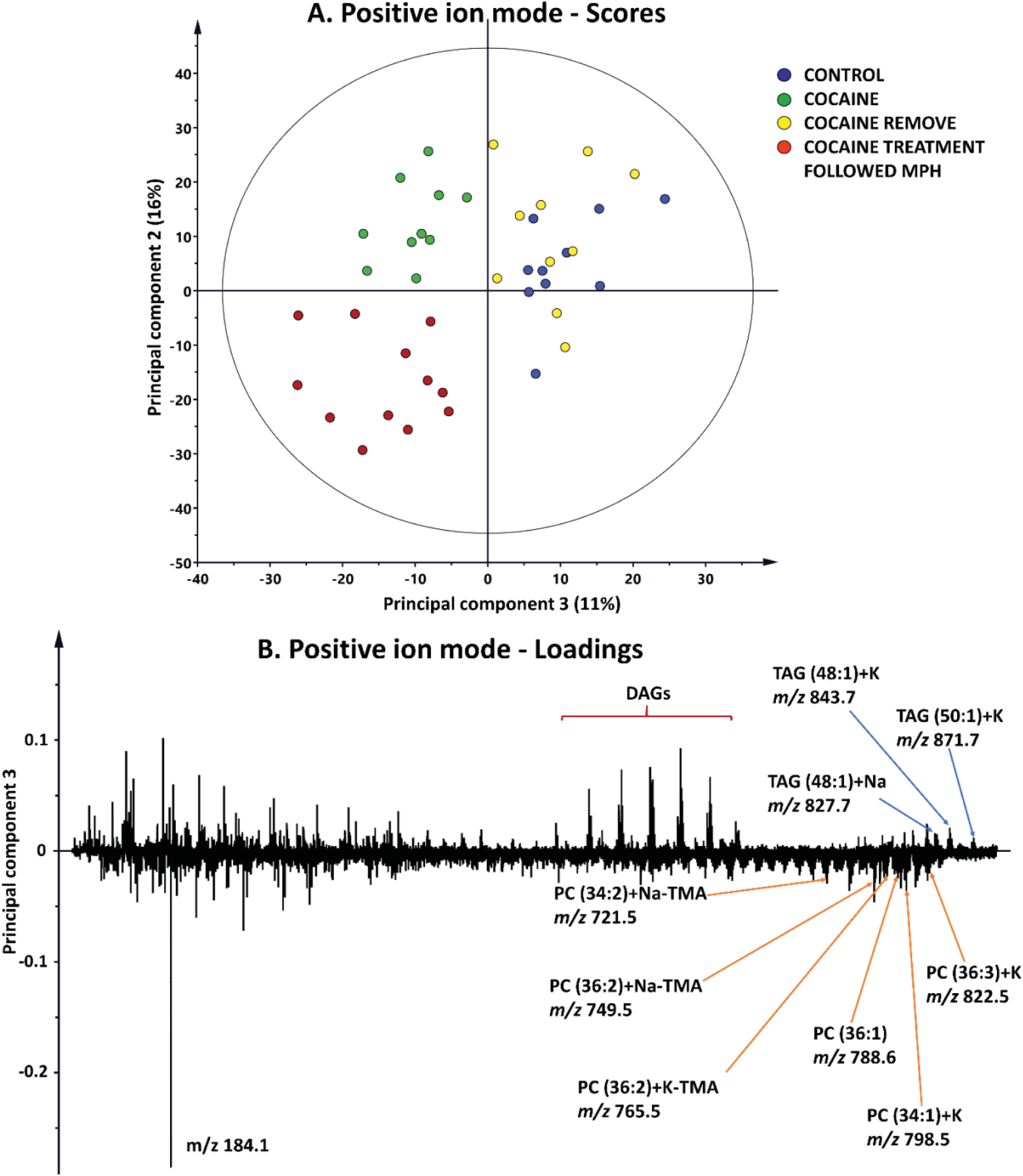
PCA for ToF-SIMS data in positive mode; (A) Score plot of the second principal component versus the third principal component; (B) Corresponding loadings for the third principal component. The ellipse on the scores plot calculated from Hotelling’s T-square statistic is related to the 95% confidence interval.

To obtain detailed information about the alterations in phospholipid compositions in fly brains after drug treatments and removal, the lipid species identified by PCA were selected and their relative concentrations were compared among different groups. Fig. 3 shows that there are significant changes in PC levels after drug treatment compared to control. PCs detected as [M+H]+ and their salt adducts are increased in the fly brains after cocaine administration. The elevation of PC abundance is mainly observed in PC species with 32, 34, and 36 carbon fatty acid chains and different saturation levels, such as PC (32:1) at *m/z* 732.6, PC (32:0) at *m/z* 734.6, PC (34:2) at *m/z* 758.6, PC (34:1) at *m/z* 760.6, PC (36:2) at *m/z* 786.6, and PC (36:1) at *m/z* 788.6. After cocaine removal, the levels of these PC species in the central brain return to the same level as in the control brains. MPH treatment during the removal time after cocaine exposure, however, induces further depletion in the PC levels compared to the control brains, which is consistent with observations in our previous reports (13, 14) and the images in Fig. 1.

**Fig. 3.**
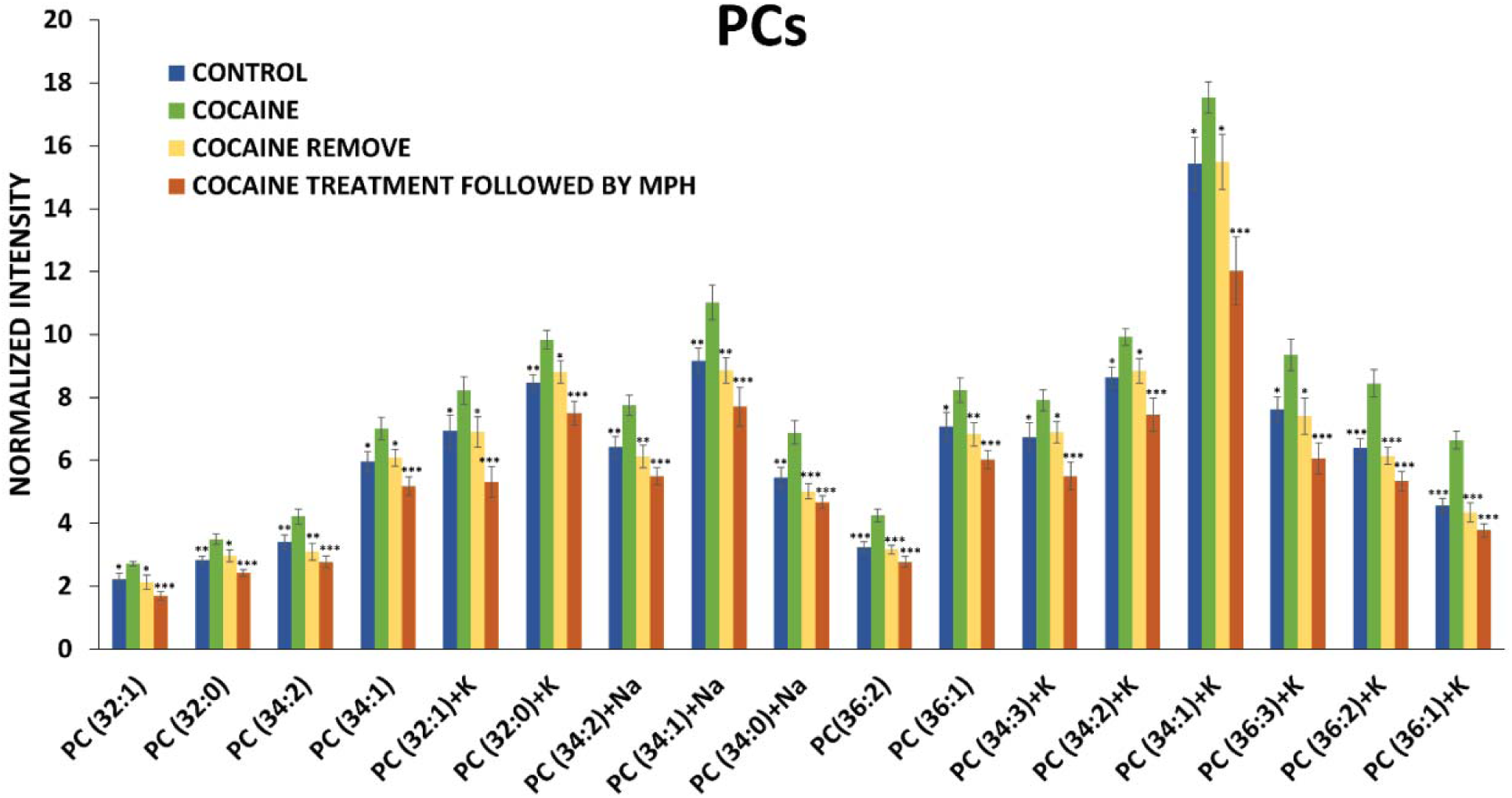
The changes in phosphatidylcholine lipid levels in the central fly brain after cocaine, cocaine removal, and flies with cocaine treatment followed MPH oral administration in positive ion mode analyzed using a 40 keV (CO_2_)_6000+_ GCIB. Peak intensity is normalized to number of pixels selected and total signal intensity. Error bars are standard error of the mean of 33 controls (blue bars), 28 cocaine-treated flies (green bars), 27 flies with cocaine withdrawal (yellow bars), and 27 flies with MPH feeding after cocaine withdrawal (red bars). A *t-test* was used to compare the statistical difference between the cocaine group with the other groups (*p<0.5, **p<0.01, ***p<0.001). No statistical differences were found between control and cocaine-removed groups (p>0.05). Ions were detected as [M+H]+ unless specified as Na/K adduct species.

The relative amounts of TAGs in different fly brain groups are shown in Fig. S2. Cocaine induces a dramatic decrease in TAG levels, and these recover partially after cocaine removal. MPH treatment after cocaine exposure and removal leads to recovery of the TAG levels to an even greater extent, but still lower than those in the control. Based on these data, we conclude that cocaine removal leads to recovery of the effects of cocaine on the abundance of PCs, and MPH further reverses the effects of cocaine on these phospholipids to an extent past control values. In addition, both treatment strategies can also partially rescue the change in TAG concentrations in fly brains following cocaine exposure.

### Changes in PE and PI induced by cocaine are not reversed by drug withdrawal

PCA was carried out on ToF-SIMS spectra in the negative ion mode to elucidate the difference for the recovery of PEs and PIs between the control and drug-treated groups. The PCA scores plot of principal component 1 versus principal component 2, which together capture 58% of the total variance in the data set, is shown in Fig. 4. The control group is clearly separated from the others across principal component 2, whereas all the drug treated groups, including the cocaine exposure, cocaine removal, and cocaine exposure followed by MPH groups, overlap in the scores plot (Fig. 4A).

**Fig. 4.**
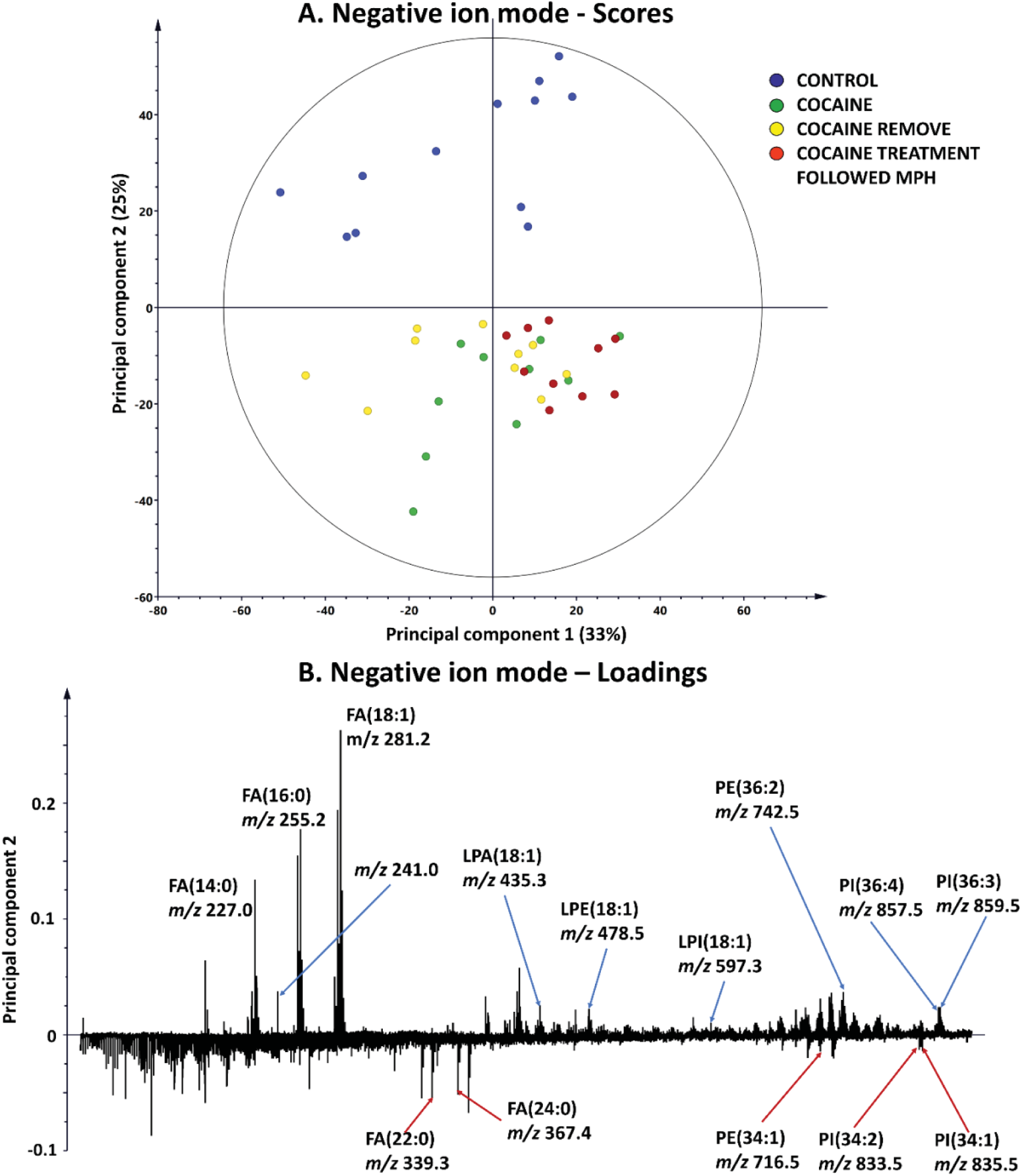
PCA for ToF-SIMS data in negative mode; (A) Score plot of the first principal component versus the second principal component; (B) Corresponding loadings for the second principal component. The ellipse on the scores plot calculated from Hotelling’s T-square statistic is related to the 95% confidence interval.

The peaks contributing to the difference between the control and the other groups in the PCA can be identified by the loadings plot of principal component 2 (Fig. 4B). The significant lipids that are decreased in the drug-treated brains versus control are PE/PI species such as PE (36:2) at *m/z* 742.5, PI (36:4) at *m/z* 857.5, and PI (36:3) at *m/z* 859.5, lysolipids such as lysophosphatidic acid LPA (18:1) at *m/z* 435.3, lysophosphatidylethanolamine LPE (18:1) at *m/z* 478.3, lysophosphatidylinositol LPI (18:1) at *m/z* 597.3, and fatty acids for example FA (18:1) at *m/z* 281.2 and FA (16:0) at *m/*z 255.2. In contrast, several PEs and PIs with 34 carbon fatty acid chain, such as PE (34:1) at *m/z* 716.5, PI (34:2) at *m/z* 833.5, and PI (34:1) at *m/z* 835.5, as well as fatty acids FA (22:0) at *m/z* 339.3 and FA (24:0) at *m/z* 367.4 are increased in the drug-treated groups.

After cocaine administration, there is an accumulation of PEs and PIs with 34 carbons in the fatty acid chain and a reduction of PEs and PIs with 36 carbons as well as several LPA, LPE and LPI species (Fig. S3). PCA, however, shows no separation between the cocaine administration and cocaine removal groups. The effect of cocaine on PEs, PIs phospholipids and several lipid precursors in the fly brain is still persistent even after the flies are removed from the drug. This is also supported by the comparison of the relative abundances of various significant lipid species in the fly brains undergoing different drug treatments (Fig 5). There is a significant increase in PEs and PIs with 34 carbon fatty acid chains, including PE (34:2), PE (34:1), PI (34:2), PI (34:1), and PI (34:0), after cocaine administration and persisting after cocaine removal. In contrast, cocaine exposure decreases the amount of several other lipids, for instance PE (30:1) at *m/z* 660.5, PE (32:1) at *m/z* 688.5, PE (36:3) at *m/z* 740.5, PE (36:2) at *m/z* 742.5, PI (36:5) at *m/z* 855.5, PI (36:4) at *m/z* 857.5, and PI (36:3) at *m/z* 859.5, and, again, the withdrawal of cocaine does not recover the effect of cocaine on these lipids. Overall, cocaine significantly alters the levels of PEs and PIs and these changes are not reversible by cocaine removal, at least for 3 days.

**Fig. 5.**
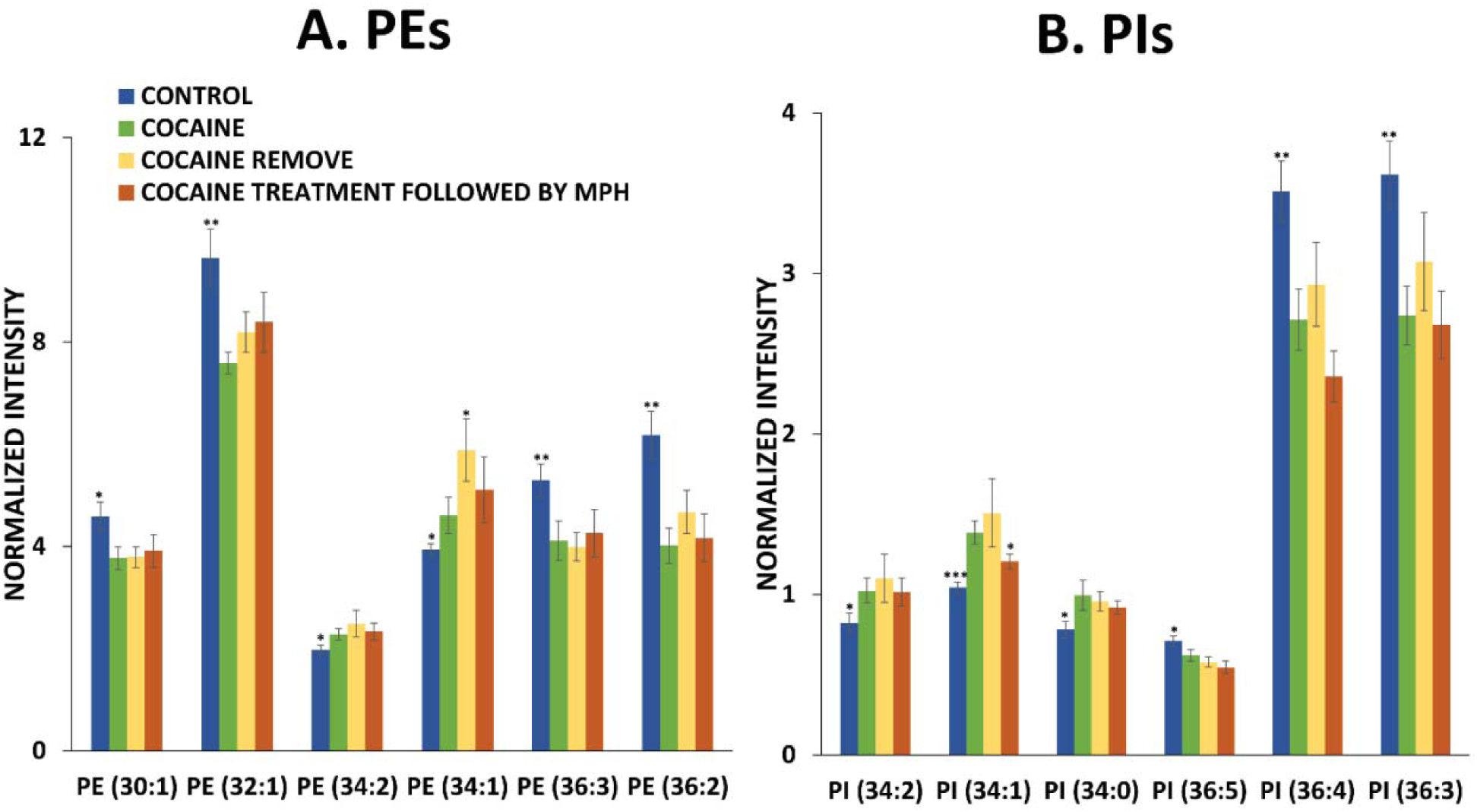
The relative amounts of phospholipids in the central fly brain after cocaine, cocaine removal, and flies with cocaine treatment followed MPH oral administration in negative ion mode analyzed using a 40 keV (CO_2_)_6000+_ GCIB. (A) PE levels; (B) PI levels. Peak intensity is normalized to number of pixels selected and total signal intensity. Error bars are standard error of the mean 32 controls (blue bars), 24 cocaine-treated flies (green bars), and 26 flies with cocaine withdrawal (yellow bars), and 24 flies fed with cocaine followed by MPH. A *t-test* was used to compare the difference between the cocaine group and the other groups (*p<0.5, **p<0.01, ***p<0.001). There are no statistical differences between cocaine groups and the cocaine-removed groups and cocaine-treated followed by MPH group (p>0.05). All ions were detected as [M-H]-.

### Changes in PE and PI induced by cocaine are not rescued by MPH treatment after cocaine removal

We also considered the effects of MPH on phospholipids in the fly brain previously exposed to cocaine, and these data are shown in Fig. 4A. Here we observe no significant difference on the brain lipid composition between the cocaine-fed flies and the cocaine-fed flies followed by MPH treatment. Fig. 5 reveals the alterations of relative levels of PEs and PIs, induced by cocaine administration to the flies compared to the control. As discussed above, cocaine causes a dramatic decrease in the concentration of most PE and PI species evaluated in the fly brain, except for the 34-carbon chain PEs and PIs, which are significantly elevated after cocaine. In contrast to the PC lipids, the levels of the PEs and PIs in the MPH treated fly brains generally stay at the same level even when they are subsequently treated with MPH for 3 days. The exception to this is that compared to the cocaine treated brains there is a slight reversal in the change of PI (34:1) treated with MPH during cocaine removal (Fig. 5). Overall, it appears that even after subsequent treatment with MPH the effects of cocaine on the composition of PE and PI lipids in the fly brain are generally persistent.

## Discussion

Oral administration cocaine results in significant changes in the phospholipid composition of the fly brain, consistent with previous results (14). PCs and the 34-carbon chain PEs and PIs are increased, and other PEs and PIs are decreased after cocaine exposure. Cocaine administration has been shown to decrease the activity of phospholipase A2 (PLA2) in the rat brain striatum due to the increase in dopamine levels (20). A schematic of the metabolic pathways of phospholipids is provided in Fig. S4. PLA2 is the enzyme responsible for the degradation of phospholipids at the sn-2 position to release 1-acyl lysophospholipids and free fatty acids (21). Reduced PLA2 activity caused by cocaine might be associated with the elevation of phospholipids, mainly PCs and some PEs and PIs, observed. In addition, the inhibition of PLA2 induced by psychostimulants such as cocaine might result in the depletion of lysophospholipids in the fly brain as it is observed in the reduction of LPA (18:1), LPE (18:1), and LPI (18:1) in the cocaine-treated flies (Fig. 4B). However, the concentrations of PCs in the fly brain return to normal values after the flies were removed from cocaine. Thus, if the PLA2 enzyme is responsible for these increased lipid levels, then it is reversible and its activity is recovered after cocaine removal (20).

Phospholipase C (PLC) is a membrane-associated enzyme that promotes the conversion of PIs to DAGs. Cocaine has been found to stimulate PLC activity via the activation of Gq/11-coupled receptors (22). PLC activity has also been found to be increased in the frontal cortex of rats after withdrawal from repeated administration of cocaine (23, 24). As the activity of PLC increases, PIs are degraded into diacylglycerol and secondary messenger molecules such as inositol 1,4,5-trisphosphate. Hence, we propose that the cocaine has long-term effects on the activity of PLC, which in turn affects the phospholipids in the long term even after withdrawal. Importantly, these persistent alterations on PEs and PIs could be associated with the cognitive deficits observed long after cocaine withdrawal. Several reports have shown that dysregulation in cognition and emotion is persistent in subjects even after cocaine withdrawal, and also after other stimulants (4, 25, 26). Since our data show that the depletion of PEs and PIs remain after cocaine removal, these changes might be associated with the cognitive impairment caused by cocaine.

An important goal for pharmacotherapeutic medicine of drug abuse is a therapeutic treatment for cocaine addiction. Several studies have suggested that MPH can be used as an agonistic substitute for treatment of cocaine dependence owing to their similarity in structure and basic mechanism in blocking dopamine uptake. However, although cocaine and methylphenidate have similar effects in blocking dopamine reuptake, they have opposite effects on their changing the phospholipid composition in fly brain (14). Moreover, it has been reported by Calipari et al. that cocaine and methylphenidate have opposite effects on the dopaminergic system (27). Cocaine decreased dopamine transport, dopamine uptake rate, and increased the dopamine release whereas MPH administration increased dopamine transporter density, dopamine uptake rate, and dopamine release. Thus, we examined the effects of MPH treatment on the phospholipid compositions of cocaine-fed flies. The feeding dose of 10 mM MPH to flies *in vivo* corresponds to a dose of 4.8 mg/kg which is in the range used for a recent pharmacokinetics and bioavailability study in rats (0.75-10 mg/kg) (28, 29). Our data (Fig. 3) reveal that PC abundance in the fly brain has a tendency to increase after cocaine administration; however, there is a downward trend or reversal when cocaine is removed, and this downward trend is exaggerated when the cocaine-fed flies are subsequently treated with MPH. In contrast, the changes in PEs and PIs in the fly brain after cocaine exposure are not reversed with cocaine removal or MPH treatment. Zhu et al. found that cocaine and MPH cause opposite effects on the release fraction during exocytosis from PC12 cells (30). They suggested that cocaine stimulated protein kinase C, which then synthesized filamentous actin and consequently increased the rate of the closing of the fusion pore during open and closed exocytosis. Unlike cocaine, MPH, which shows no effects on actin immunoactivity (31), shows normal exocytosis similar to control cells. Thus, a possible mechanistic aspect to the effects of cocaine and MPH might be that they induce various effects on membrane phospholipids, which then link to the stability of the fusion pores for exocytosis.

We observe that PC levels are increased in the central fly brain following cocaine exposure. An elevation of PC abundance was also observed in rat brains treated with other stimulants which also induces the impairment of cognitive performance (32). There are now several pieces of evidence suggesting that brain phospholipids, especially PCs, are involved in brain function and mental performance (33, 34). PCs are closely related to the production of choline which is a precursor for acetylcholine, a neurotransmitter that plays many biological roles in developmental and cognitive function (35). PC levels decrease again to normal levels in fly brains after cocaine removal, or after MPH treatment. In addition to ADHD patients, cognitive enhancement following MPH has also been reported for patients with traumatic brain injury as well as depression after stroke (36–38). Moreover, Goldstein’s group showed that the use of a low dose of MPH improved cognitive tasks in cocaine abusers (39). It is enticing to speculate that the changes we observe in lipids following cocaine and those rescued by MPH are in some way involved in these cognitive changes. It should be noted that incubation of cells with lipids changes the rate of exocytosis (40, 41).

In summary, PCs, PEs, PIs might be considered to be therapeutic targets for treatment of cognitive deficit caused by cocaine. Although cocaine removal and MPH treatment after cocaine removal do not completely reverse the dramatic alterations of all phospholipids in the brain caused by cocaine, they impose a significantly positive effect on the adverse action of cocaine on PCs and partially on TAGs. The approaches presented here might be further examined to develop rescue strategies for the cognitive impairment by cocaine addiction.

## Acknowledgments

This study was supported by the Knut and Alice Wallenberg Foundation, the European Research Council (ERC) and the Swedish Research Council (VR).

## SUPPLEMENTAL INFORMATION

**Supplemental Table S1.** Assignment for selected peaks from fly brain in positive and negative ion modes.

| | Exact m/z | Measured m/z | Δppm | Label | Species |
| --- | --- | --- | --- | --- | --- |
| **Positive ion mode** | 713.4482 | 713.4501 | 2.7 | PC(32:0) | [M+K-TMA]<sup>+</sup> |
|  | 721.4779 | 721.4802 | 3.2 | PC(34:2) | [M+Na-TMA]<sup>+</sup> |
|  | 732.5560 | 732.5485 | -2.0 | PC(32:1) | [M+H]<sup>+</sup> |
|  | 734.5655 | 734.5621 | -4.6 | PC(32:0) | [M+H]<sup>+</sup> |
|  | 739.4714 | 739.4668 | -6.2 | PC(34:1) | [M+K-TMA]<sup>+</sup> |
|  | 749.5135 | 749.5154 | 2.5 | PC(36:2) | [M+Na-TMA]<sup>+</sup> |
|  | 758.5652 | 758.5626 | -3.4 | PC(34:2) | [M+H]<sup>+</sup> |
|  | 760.5846 | 760.5809 | -4.9 | PC(34:1) | [M+H]<sup>+</sup> |
|  | 765.4753 | 765.4693 | -7.8 | PC(36:2) | [M+K-TMA]<sup>+</sup> |
|  | 770.5129 | 770.5096 | -4.3 | PC(32:1) | [M+K]<sup>+</sup> |
|  | 772.5243 | 772.5218 | -3.2 | PC(32:0) | [M+K]<sup>+</sup> |
|  | 780.5514 | 780.5464 | -6.4 | PC(34:2) | [M+Na]<sup>+</sup> |
|  | 782.5612 | 782.5563 | -6.3 | PC(34:1) | [M+Na]<sup>+</sup> |
|  | 784.5606 | 784.5568 | -4.8 | PC(34:0) | [M+Na]<sup>+</sup> |
|  | 786.5809 | 786.5768 | -5.2 | PC(36:2) | [M+H]<sup>+</sup> |
|  | 788.6209 | 788.6139 | -5.9 | PC(36:1) | [M+H]<sup>+</sup> |
|  | 794.5063 | 794.5024 | -4.9 | PC(34:3) | [M+K]<sup>+</sup> |
|  | 796.5218 | 796.5186 | -4.0 | PC(34:2) | [M+K]<sup>+</sup> |
|  | 798.5396 | 798.5474 | -2.7 | PC(34:1) | [M+K]<sup>+</sup> |
|  | 822.5410 | 822.5398 | -1.4 | PC(36:3) | [M+K]<sup>+</sup> |
|  | 824.5513 | 824.5487 | -3.1 | PC(36:2) | [M+K]<sup>+</sup> |
|  | 826.5702 | 826.5687 | -1.8 | PC(36:1) | [M+K]<sup>+</sup> |
|  | 827.7099 | 827.7054 | -5.4 | TAG(48:1) | [M+Na]<sup>+</sup> |
|  | 841.7256 | 841.7208 | -5.7 | TAG(48:2) | [M+K]<sup>+</sup> |
|  | 843.7412 | 843.7439 | 3.2 | TAG(48:1) | [M+K]<sup>+</sup> |
|  | 853.7256 | 853.7208 | -5.6 | TAG(50:2) | [M+Na]<sup>+</sup> |
|  | 855.7412 | 855.7358 | -6.3 | TAG(50:1) | [M+Na]<sup>+</sup> |
|  | 871.7225 | 871.7186 | -4.5 | TAG(50:1) | [M+K]<sup>+</sup> |
|  | 879.7412 | 879.7376 | -4.1 | TAG(52:3) | [M+Na]<sup>+</sup> |
|  | 881.7569 | 881.7535 | -3.8 | TAG(52:2) | [M+Na]<sup>+</sup> |
|  | 883.7725 | 883.7697 | -3.2 | TAG(52:1) | [M+Na]<sup>+</sup> |
| **Negative ion mode** | 253.2173 | 253.2185 | 4.7 | FA(16:1) | [M-H]<sup>-</sup> |
|  | 255.2330 | 255.2347 | 6.7 | FA(16:0) | [M-H]<sup>-</sup> |
|  | 279.2330 | 279.2346 | 5.7 | FA(18:2) | [M-H]<sup>-</sup> |
|  | 281.2486 | 281.2497 | 3.9 | FA(18:1) | [M-H]<sup>-</sup> |
|  | 283.2643 | 283.2657 | 4.9 | FA(18:0) | [M-H]<sup>-</sup> |
|  | 339.3269 | 339.3287 | 5.3 | FA(22:0) | [M-H]<sup>-</sup> |
|  | 367.3582 | 367.3602 | 5.4 | FA(24:0) | [M-H]<sup>-</sup> |
|  | 435.2517 | 435.2533 | 3.7 | LPA(18:1) | [M-H]<sup>-</sup> |
|  | 478.2939 | 478.2965 | 5.4 | LPE(18:1) | [M-H]<sup>-</sup> |
|  | 597.3045 | 597.3081 | 6.0 | LPI(18:1) | [M-H]<sup>-</sup> |
|  | 660.4567 | 660.4534 | -5.0 | PE(30:1) | [M-H]<sup>-</sup> |
|  | 688.4878 | 688.4831 | -6.8 | PE(32:1) | [M-H]<sup>-</sup> |
|  | 714.5053 | 714.4998 | -7.7 | PE(34:2) | [M-H]- |
|  | 716.5223 | 716.5201 | -3.1 | PE(34:1) | [M-H]- |
|  | 740.5236 | 740.5195 | -5.5 | PE(36:3) | [M-H]- |
|  | 742.5382 | 742.5334 | -6.5 | PE(36:2) | [M-H]- |
|  | 833.5149 | 833.5125 | -2.9 | PI(34:2) | [M-H]- |
|  | 835.5298 | 835.5298 | -6.0 | PI(34:1) | [M-H]- |
|  | 837.6070 | 837.5989 | -2.4 | PI(34:0) | [M-H]- |
|  | 855.5041 | 855.5015 | -3.0 | PI(36:5) | [M-H]- |
|  | 857.5148 | 857.5098 | -5.8 | PI(36:4) | [M-H]- |
|  | 859.5342 | 859.5295 | -5.5 | PI(36:3) | [M-H]- |

**Supplemental Fig. S1.**
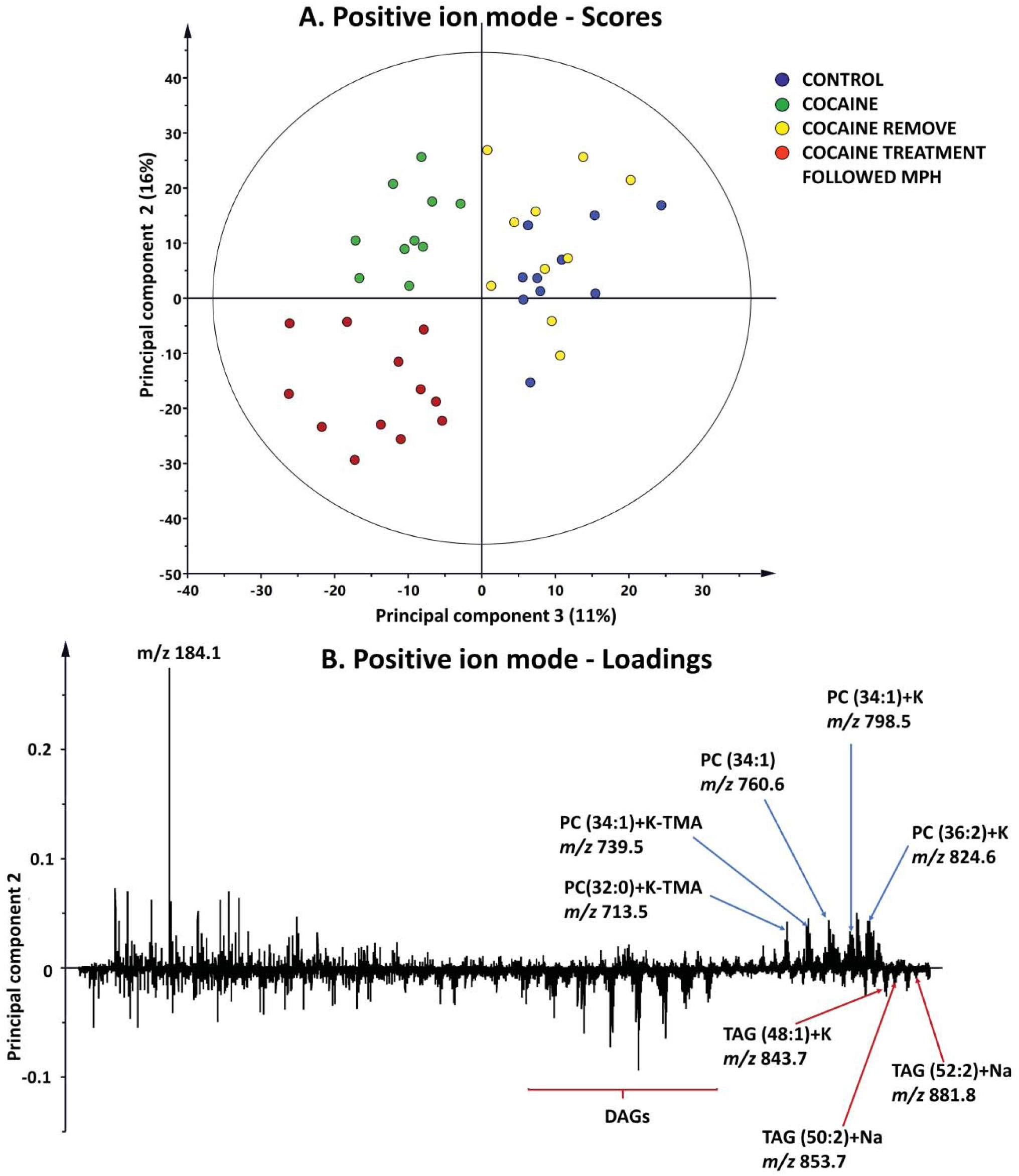
PCA analysis for ToF-SIMS data in positive mode; (A) Scores plot of the second principal component versus the third principal component; (B) Corresponding loadings for the second principal component for ToF-SIMS data in positive mode.

**Supplemental Fig. S2.**
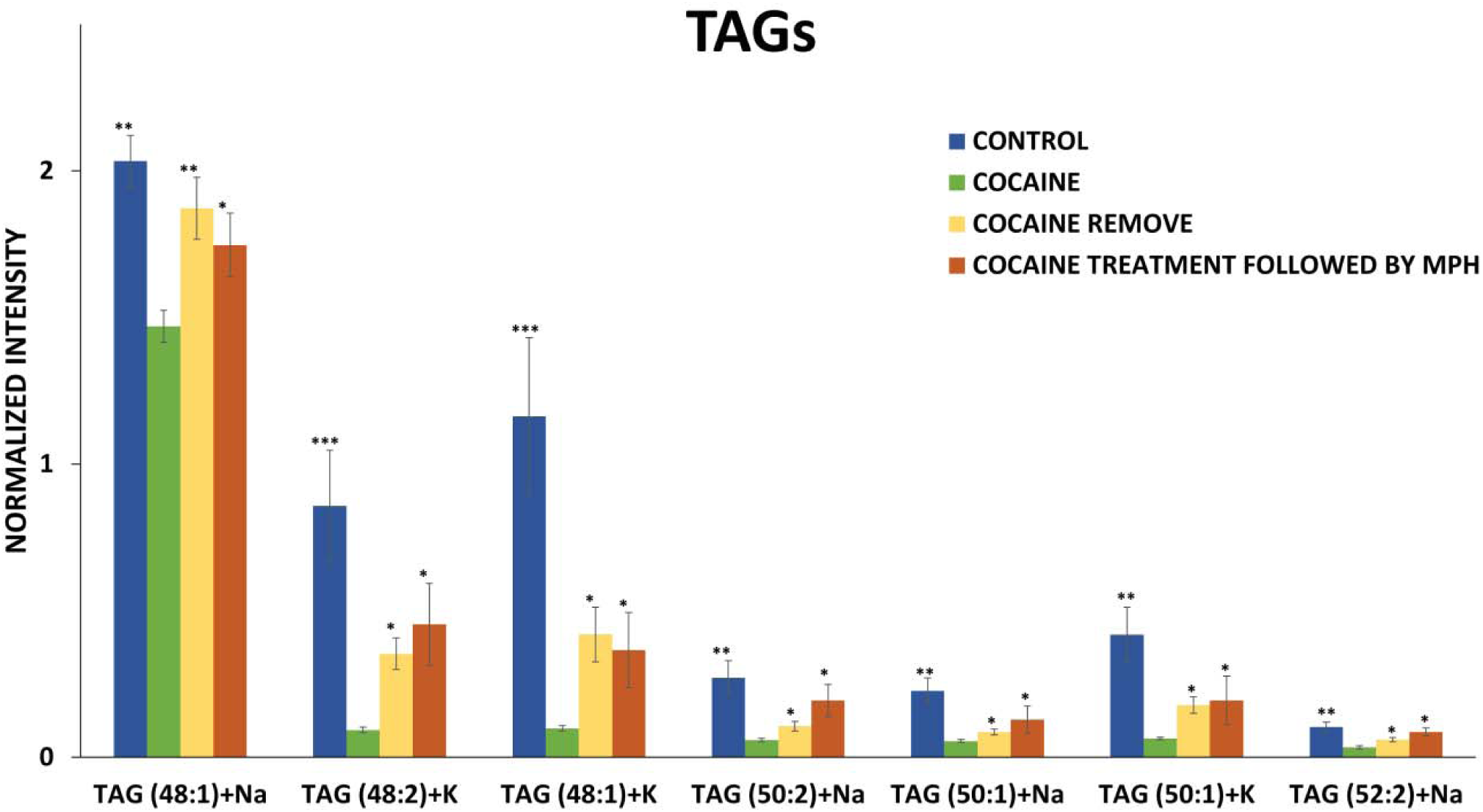
Alteration of TAG levels in the central fly brain after cocaine, cocaine removal, and flies with cocaine treatment followed MPH oral administration in positive ion mode analyzed using a 40 keV (CO_2_)_6000+_ GCIB. Peak intensity is normalized to number of pixels selected and total peak intensity. Error bars is the standard error of the mean of 33 controls (blue bars), 28 cocaine-treated flies (green bars), 27 flies with cocaine withdrawal (yellow bars), and 27 flies with MPH feeding after cocaine withdrawal (red bars). *t-test* is applied to identify the statistic difference between cocaine-treated group and others (*p<0.05, **p<0.01, ***p<0.001).

**Supplemental Fig. S3.**
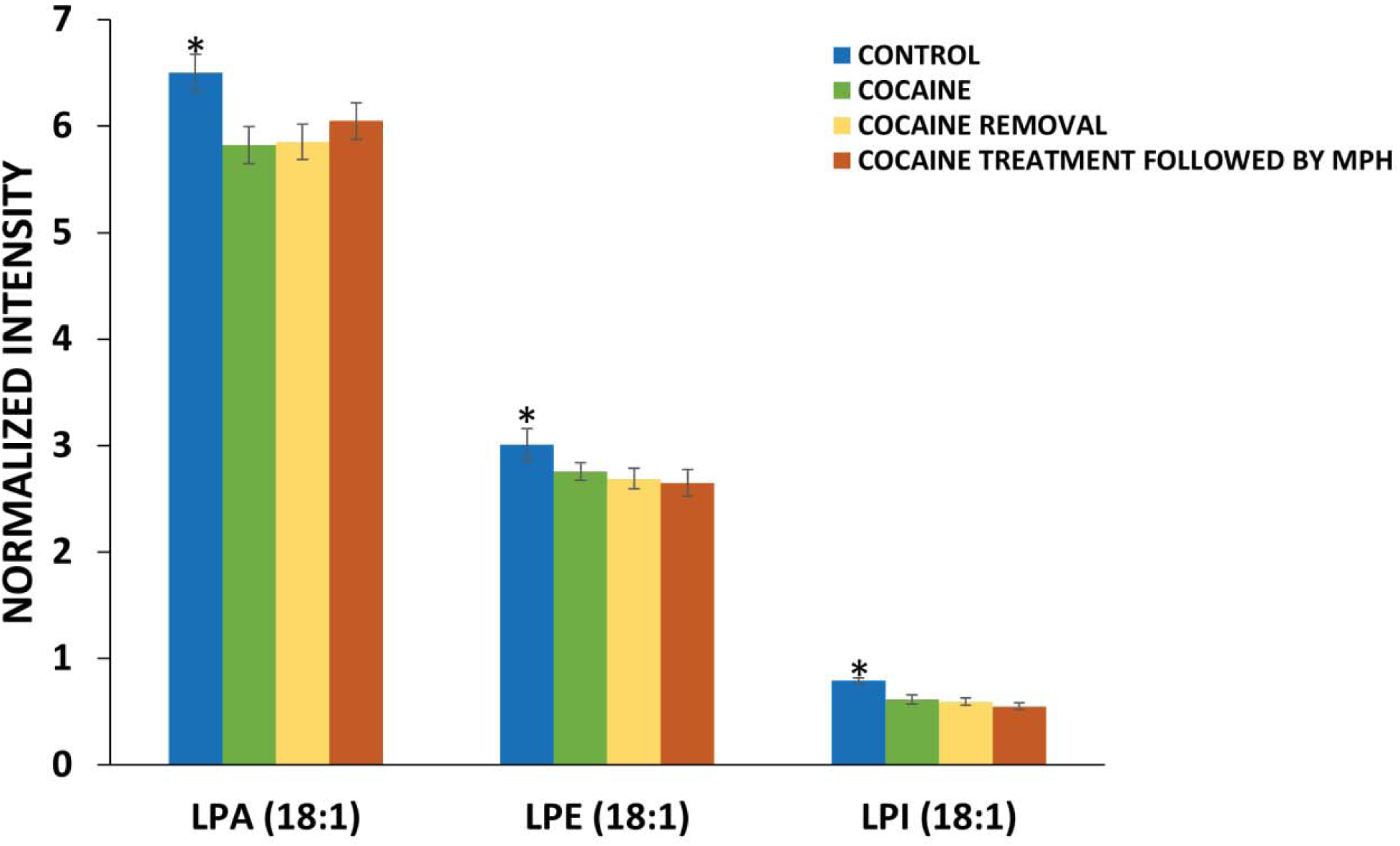
The relative amounts of lysophospholipids in the central fly brain after cocaine, cocaine removal, and flies with cocaine treatment followed MPH oral administration in negative ion mode analyzed using a 40 keV (CO_2_)_6000+_ GCIB. (A) PE levels; (B) PI levels. Peak intensity is normalized to number of pixels selected and total peak intensity. Error bars is the standard error of the mean of 32 controls (blue bars), 24 cocaine-treated flies (green bars), and 26 flies with cocaine withdrawal (yellow bars), and 24 flies fed with cocaine followed by MPH. *t-test* is applied to identify the statistic difference between cocaine-treated group and others (*p<0.05). There is no statistic difference between cocaine group and cocaine removal group, as well as cocaine-treated followed by MPH group (p>0.05).

**Supplemental Fig. S4.**
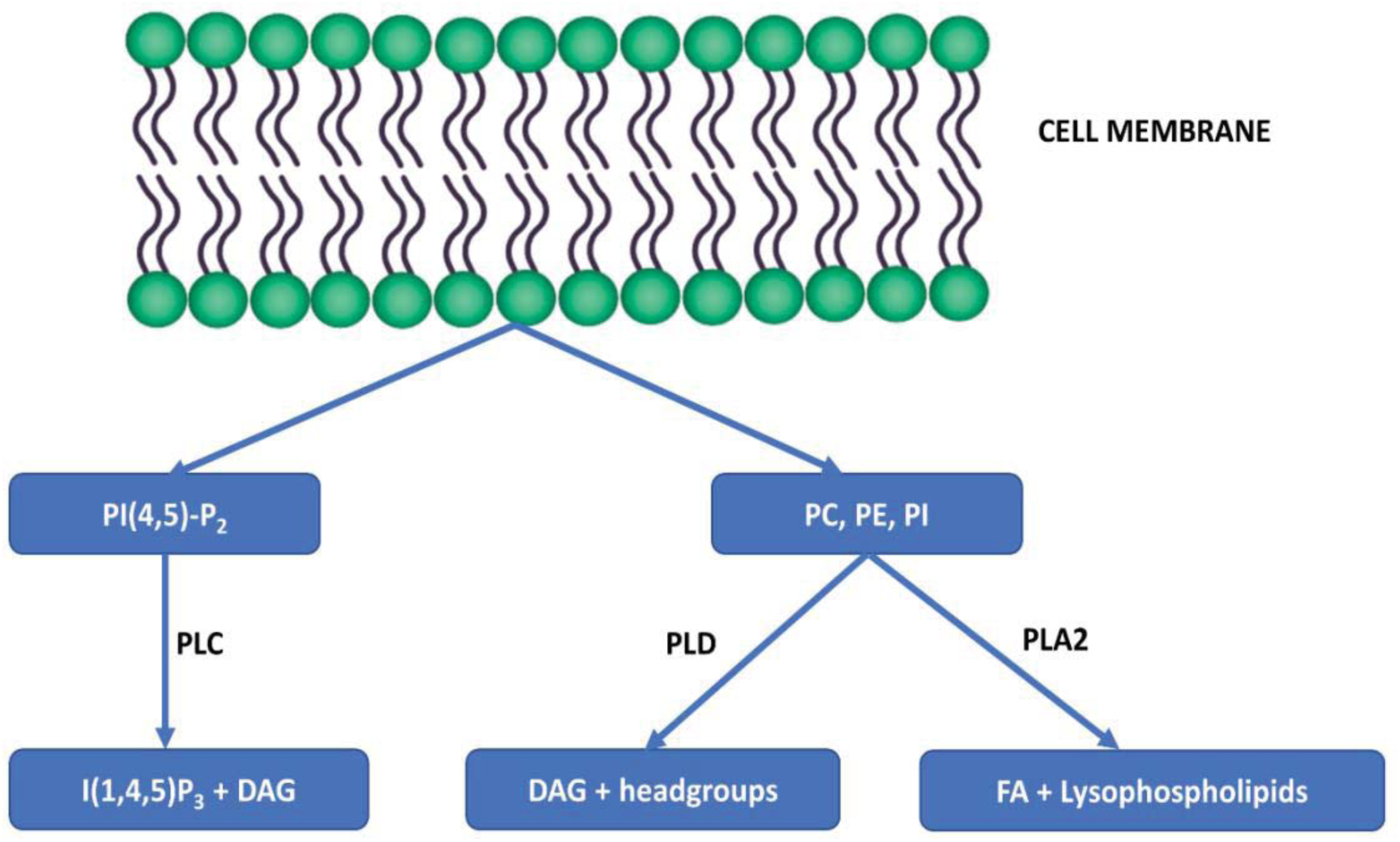
Scheme showing the degradation of membrane phospholipids by phospholipase A2, C, and D (PLA2, PLC, and PLD). PI (4,5)P2, phosphatidylinositol 4,5-biphosphate; IP3, inositol 1,4,5-triphosphate; DAG, diacylglycerol; PC, phosphatidylcholine; PE, phosphatidylethanolamine; PI, phosphatidylinositol.

